# The impact of pubertal stress and adult hormone exposure on the transcriptome of the developing hypothalamus

**DOI:** 10.1101/2023.10.03.559350

**Authors:** Karissa N. Gautier, Samantha L. Higley, John M. Mendoza, Kathleen E. Morrison

## Abstract

Why individuals suffer negative consequences following stress is a complex phenomenon that is dictated by individual factors, the timing of stress within the lifespan, and when in the lifespan the consequences are measured. Women who undergo adverse childhood experiences are at risk for lasting biological consequences, including affective and stress dysregulation. We have shown that pubertal adversity is associated with a blunted hypothalamic-pituitary-adrenal axis glucocorticoid response in peripartum humans and mice. In mice, our prior examination of the paraventricular nucleus (PVN) of the hypothalamus showed that pubertal stress led to an upregulation of baseline mRNA expression of six immediate early genes (IEGs) in the PVN of adult, pregnant mice. Separately, we showed that the pregnancy-associated hormone allopregnanolone is necessary and sufficient to produce the blunted stress response phenotype in pubertally stressed mice. In the current study, we further examined a potential mechanistic role for the IEGs in the PVN. We found that in pubertally stressed adult female, but not male, mice, intra-PVN allopregnanolone was sufficient to recapitulate the baseline IEG mRNA expression profile previously observed in pubertally stressed, pregnant mice. We also examined baseline IEG mRNA expression during adolescence, where we found that IEGs have developmental trajectories that showed sex-specific disruption by pubertal stress. Altogether, these data establish that IEGs may act as a key molecular switch involved in increased vulnerability to negative outcomes in adult, pubertally stressed animals. How the factors that produce vulnerability combine throughout the lifespan is key to our understanding of the etiology of stress-related disorders.

## Introduction

Risk for neuropsychiatric disorders in adulthood is increased by exposure to adverse childhood experiences (ACEs), which are events including emotional abuse, physical abuse, and neglect that happen up to the age of 18 (1,2). Women exposed to ACEs during puberty are at the greatest risk for neuropsychiatric disorders across the lifespan (3–5) and suffer unique physiological and biological symptoms compared to those who experienced adversity earlier in childhood or adulthood (6–8). We previously established that chronic stress during puberty in female mice mimicked pubertal ACEs in women, where mice and humans had a significantly blunted stress reactivity in adulthood, but only during the peripartum window (9). Risk of peripartum depression and anxiety are elevated in women with high lifetime adversity, and perinatal mood disorders are associated with long-term negative outcomes for both mother and baby (10). Although we know that multiple experiences (e.g., pubertal stress, pregnancy) compound over the lifespan to influence the risk for negative outcomes, we have limited understanding of how this risk manifests at the biological level.

We have previously shown lasting effects of stress during puberty on the physiology and behavior of female mice and humans (8,9,11,12). Mice and humans exposed to pubertal stress showed a blunted hormonal stress response (Figure 1), which is associated with increased postnatal depression scores in humans. To understand the mechanisms underlying these lasting effects of pubertal stress, our focus has been on the paraventricular nucleus of the hypothalamus (PVN), a key brain region that regulates the hypothalamic-pituitary-adrenal (HPA) stress axis response and a region we have previously identified to be disrupted in both the epigenome and transcriptome following pubertal stress in mice (9,11). Transcriptomic analysis of the PVN during late pregnancy identified six immediate early genes (IEGs) that were permissively expressed at baseline in the PVN of pregnant, pubertally-stressed females. Here, we leveraged this pubertal stress-associated transcriptional signature to provide insight into the molecular underpinnings of the lasting effects of early life stress or trauma, a phenomenon not well understood (13–15).

**Figure 1.**
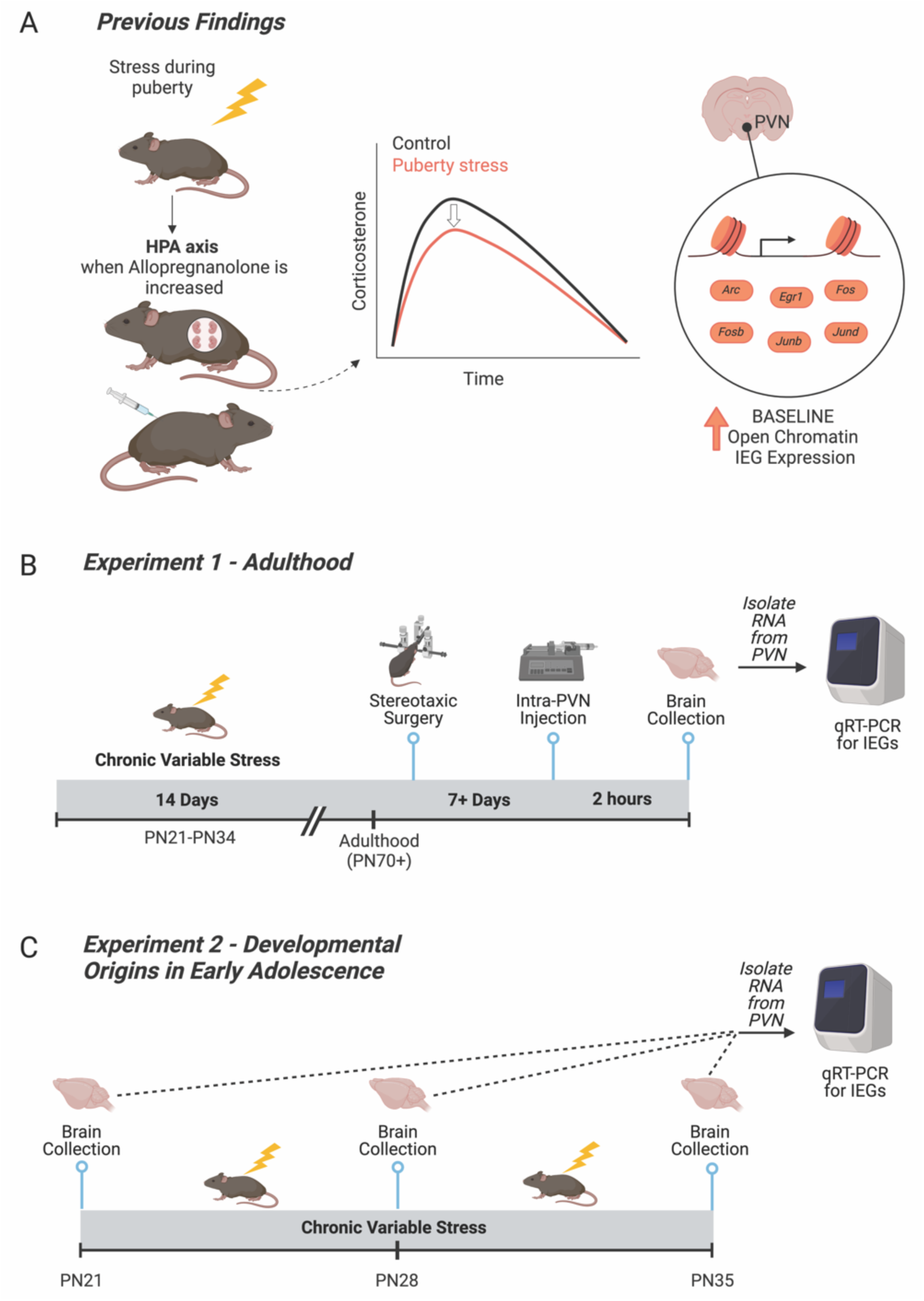
Relevant prior findings and experimental timelines for the current study. **(A)** We previously established that chronic variable stress (CVS) during puberty leads to a blunted hypothalamic-pituitary-adrenal axis in adulthood only when allopregnanolone is present at high levels in the brain, including during pregnancy and under allopregnanolone treatment (9,11). We identified the paraventricular nucleus of the hypothalamus (PVN) as a key tissue of the HPA axis that was reprogrammed by pubertal stress. Pubertal stress and pregnancy combined to lead to an altered chromatin landscape, including an increase in the number of open chromatin sites, and permissive expression of six immediate early genes (IEGs) at baseline. These findings suggest that IEGs play a mechanistic role in the puberty-stress induced phenotype that is apparent during times of dynamic change in allopregnanolone, including pregnancy. **(B)** Here, we directly tested the relationship between allopregnanolone within the PVN and the puberty stress associated IEG expression. Female and male mice were exposed to CVS from PN21-34. In adulthood, mice were implanted with a double-barrel cannula aimed at the PVN. Following recovery, mice were injected with either vehicle or allopregnanolone. Two hours later, brains were collected and processed for the measurement of gene expression. **(C)** We further examined a potential mechanistic role for IEG expression during the pubertal stress window. Brains were collected in baseline, non-stimulated conditions from female and male mice at either at PN21 (Control only), PN28 (Control and CVS, 24 hours after 1 week of pubertal stress), or PN35 (Control and CVS, 24 hours after the full 2 weeks of pubertal stress). Brains from CVS mice were collected 24 hours after exposure to the last stressor. Created with BioRender.com.

We have also previously shown that neurosteroid allopregnanolone may be the aspect of pregnancy that is responsible for unmasking the disrupted HPA axis response in pubertally stressed mice, as it is both necessary and sufficient to block and recapitulate the blunted HPA axis, respectively (11). The function of allopregnanolone as an FDA-approved treatment for postpartum depression (Brexanolone, Zuranolone) is attributed to its action at the GABA-A receptor (16–18). While the rise in allopregnanolone that occurs during pregnancy initiates important plasticity in GABA-A receptors, as well as regulates the HPA axis and oxytocin systems in the brain (19–21), in our studies allopregnanolone also increased risk for negative outcomes in perinatal, pubertally stressed mice. One potentially meaningful difference between the models that established allopregnanolone as an effective postpartum depression treatment is that those do not involve early life stress and our model does.

Whether allopregnanolone and the IEGs are linked in producing this increased risk is unknown. Although IEG protein expression is associated with neural activation and plasticity after exposure to stimuli, IEG mRNA and proteins are canonically very low to undetectable at baseline (22–24). So, the increased expression of IEG mRNA at baseline in pregnant, pubertally stressed females suggests a potential mechanistic role in influencing how the PVN is poised to respond to the environment. Here, we examined how risk for negative outcomes in females manifests at the molecular level by taking an approach that includes assessment of risk-associated changes to the transcriptome of the hypothalamus, both during the stress exposure in puberty and later in life when the phenotype appears (pregnancy).

Therefore, we examined the baseline mRNA expression of the IEGs in the PVN under relevant pharmacological conditions and at various ages. First, in adulthood we pharmacologically manipulated allopregnanolone specifically in the PVN of adult, pubertally stressed mice to address whether this treatment alone was sufficient to recapitulate the increase in IEG mRNA expression in baseline conditions (**Figure 1B**). Second, we examined the developmental origins of the increase in baseline IEG mRNA expression and whether pubertal stress influenced the stability of baseline gene expression throughout the early adolescence window (**Figure 1C**).

## Methods

### Animals

All mice were virgin, in house mixed strain C57BL/6:129 (N = 143) (9,11,12). Some mice were heterozygous *Crh-IRES-Cre;Ai14* C57BL/6:129, capable of expressing tdTomato in corticotropin releasing factor cells, although the transgenic background was not utilized in this study, nor does it produce differing outcomes from wild-types of the same strain (25,26). All mice were kept on a 12-hour light-dark cycle (lights on at 8:00 am) and food and water were available *ad libitum*. All procedures were approved by the West Virginia University Institutional Animal Care and Use Committee and were conducted in accordance with the National Institutes of Health Guide for the Care and Use of Laboratory Animals.

### Pubertal stress

Starting on postnatal day (PN) 21, mice underwent two weeks of chronic variable stress (CVS) (9,11,12). The stressors began within the first hour of lights on (on at 8:00 am) during the daily nadir of circadian corticosterone levels. CVS lasted two hours each day, excluding the acute restraint, which was only 15 minutes in length. Two stressors were randomly assigned to be applied simultaneously each day and varied between three different sensory modalities: tactile (wet bedding, wire mesh, no bedding, toy glass marbles, multiple cage changes (3x), 15-minute acute restraint stress), auditory (white noise, owl screech), and olfactory (70% ethanol; puma odor [1:200 2-Phenethylamine, CAS 64-04-0, in mineral oil]). Mice experienced rotating combinations of stressors (tactile and auditory, tactile and olfactory, or auditory and olfactory), combinations of which also rotated between specific stressors in each category. For Experiment 1, mice in the CVS group were weaned into singly housed cages at the beginning of stress (PN21) and were pair-housed with a same-sex, same-stress littermate at the end of the 14 days of stress (PN34). Control mice were weaned on PN28 into same-sex, pair-housed cages. Mice in the CVS group were weaned at PN21 to include early neglect as part of the stressor experience, consistent with our previous work in this model (9,11,12). For Experiment 1, they were left undisturbed until adulthood (18-19 weeks). For Experiment 2, mice in the CVS group were weaned into singly housed cages at the beginning of stress (PN21). They were left undisturbed for at least 24 hours after the last stressor and prior to brain collection, which occurred at PN28 or PN35. Control mice collected at PN21 and PN28 were removed directly from the litter at the time of collection. Control mice used for PN35 collections were weaned on PN28 into same-sex, pair-housed cages.

### Experiment 1: pharmacological manipulation of allopregnanolone in adulthood

We previously observed pubertal stress-associated physiological, transcriptomic, and chromatin outcomes in adulthood when outcomes were examined under conditions of high allopregnanolone (pregnancy or peripheral allopregnanolone treatment, **Figure 1A**). Here we sought to mechanistically link allopregnanolone to the transcriptomic outcomes, a phenotype where six immediate early genes (IEGs) are permissively expressed in the PVN at baseline conditions. (**Figure 1B**). To test a direct role for allopregnanolone in the IEG phenotype in the PVN, adult male and female mice were treated with allopregnanolone (100 ng per side of PVN; Tocris 3653) or vehicle (20% weight/volume 2-Hydroxypropyl-β-cyclodextrin in water, Tocris 0708). Allopregnanolone was suspended in vehicle at 100ng/200nL for intra-PVN administration. It was prepared in advance and stored at -20°C.

In adulthood (age 18-19 weeks), female (N = 26) and male (N = 19) mice underwent stereotaxic cannulation to selectively target the PVN. Mice were anesthetized using 4% isoflurane in an induction chamber and immediately connected to the surgery setup, where they received constant isoflurane (0.5%-2.5%). In addition, local analgesic carprofen (SC, 5 mg/kg) and anesthetics bupivacaine (at incision site, 1.5 mg/kg) and lidocaine (at incision site, 0.5 mg/kg) were administered before beginning the procedure. A double-barreled cannula (0.6mm center-to-center placing, 4mm cut below pedestal, P1 Technologies custom) was implanted in the brain at the PVN relative to bregma (-0.3mm medial/lateral, -0.85mm anterior/posterior) and lowered 3.8mm below the skull. Dental cement was used to fill the incision and create a headcap around the cannula, securing it to the skull. Mice were monitored for five consecutive days and received peripheral carprofen (SC, 5 mg/kg) for the first three days post-surgery. All mice were observed for 6 days prior to further manipulation. Mice were infused with allopregnanolone between 11 to 22 days later (age 21 or 23 weeks). On the day of infusion, mice were gently restrained, and injection needles were inserted that projected 2 mm past the cannula guides. 200nL of allopregnanolone (100 ng/200 nl) or 200 nl vehicle was simultaneously infused into each side of the double-barreled cannula using two Hamilton Microsyringes and Dual Syringe Nanoliter Pump over a 1 minute period. The needle was left in place for an additional 1 minute to allow the infusion to diffuse away from the needle. Mice were briefly restrained to accomplish the insertion and removal of the injection needle. The dose and timing have been shown effective in altering behavior in prior studies (27,28). Mice were euthanized two hours later via cervical dislocation. Brains were frozen on dry ice and stored at -80°C until use.

### Experiment 2: Influence of CVS on IEG expression during adolescence

Brains were collected from female and male mice at one of three time points: PN21 (prior to the start of CVS), PN28 (after 1 week of CVS), or PN35 (after 2 weeks of CVS). At PN28 and PN35, brains were either from Control or CVS mice 24 hours after the last stressor. Sample sizes started at N=10 per group, although there was some attrition (**Table S1**). Mice were anesthetized with vaporized Isoflurane and brains were collected under baseline (non-stimulated) conditions. Brains were frozen on dry ice and stored at -80°C until use.

### PVN collection and analysis

Brains were cryosectioned to collect precise samples of the PVN. Two, 300 um slices were taken from the brain at the anatomic location of the PVN using Paxinos and Franklin’s stereotaxic coordinates (29) for all brains, except those from PN21 animals, for which we utilized two, 250 um slices. Pre-chilled 1mm biopsy tissue punchers were used to remove the PVN from each slice. PVN biopsies were ejected into a clean, pre-chilled 1.5 mL tube and stored at -80°C until use.

Verification of accurate cannula placement for Experiment 1 was made by inspecting the 300um slices of each sample for clear indications that the cannula and infusion needles had passed through the tissue. Further confirmation of accurate PVN location was completed by taking 10um slices before and after the region. The sections were stained using neutral red to verify anterior and posterior sites around the PVN. Samples were omitted from the study if the injection site fell outside of this region or neutral red staining revealed incorrect cryosectioning of the PVN (1 male and 1 female, **Table S1**).

### Gene expression

We examined the expression of the six target IEGs (*Fos, Fosb, Arc, Egr1, Junb,* and *Jund)*. RNA was isolated from PVN tissue using the RNeasy Mini Kit (Qiagen 74106) following the manufacturer’s instructions. The quality of isolated RNA was assessed using a NanoDrop and Qubit RNA High Sensitivity Kit (Invitrogen Q32852). The Applied Biosystems High-Capacity cDNA Reverse Transcriptase Kit (#4368814) was used to generate cDNA, and the quality of cDNA was assessed on a NanoDrop.

Gene expression was measured using the QuantStudio 5 quantitative real-time PCR (qRT-PCR) system. All gene expression assays were prepared with TaqMan Fast Advanced Master Mix (Applied Bio 4444556) and probes for the target genes and reference gene *Gapdh* (**Table S2**). Each reaction was run in triplicate using an equal volume of cDNA and *Gapdh* was run on each plate. Relative gene expression was calculated using the ΔΔCt method.

### Statistical analyses

For experiment 1, gene expression data were analyzed by a 3-way ANOVA with sex, stress, and treatment (vehicle vs. allopregnanolone) as factors. Interactions were further investigated by 2-way ANOVAs (stress x treatment) within sex. For experiment 2, gene expression data were analyzed using several intersecting approaches: independent samples t-test for baseline sex differences at PN21, linear regression for trajectory of gene expression from PN21 to PN35, multiple regression to examine the impact of sex, stress, and age on changes in gene expression throughout early adolescence, and 3-way ANOVA with sex, stress, and age (P28, PN35) as factors. Bonferroni post-hoc tests were utilized following any 2-way ANOVA. Data were considered outliers and excluded from analysis if they were ± two standard deviations from the mean. Sample sizes for all groups are provided in the Supplement (**Table S1**). All analyses were performed in Prism (GraphPad) with an alpha level of *p* < 0.05. Detailed output of all statistical tests are provided in the Supplement.

## Results

### Intra-PVN allopregnanolone is sufficient to recapitulate the pubertal stress associated baseline IEG mRNA expression profile in adult females

We have previously shown that pubertal stress (CVS) resulted in a blunted corticosterone response to acute restraint stress in adult mice during pregnancy or with allopregnanolone treatment (**Figure 1A**). Further, we identified a distinct epigenetic phenotype in the PVN in pregnant, pubertally stressed females, such that the chromatin landscape was open and there was permissive expression of six IEGs at baseline. Here, we tested whether it is the increased levels of allopregnanolone present in pregnancy that interact with prior pubertal stress experience to increase baseline IEG expression in the PVN (**Figure 2A**).

**Figure 2.**
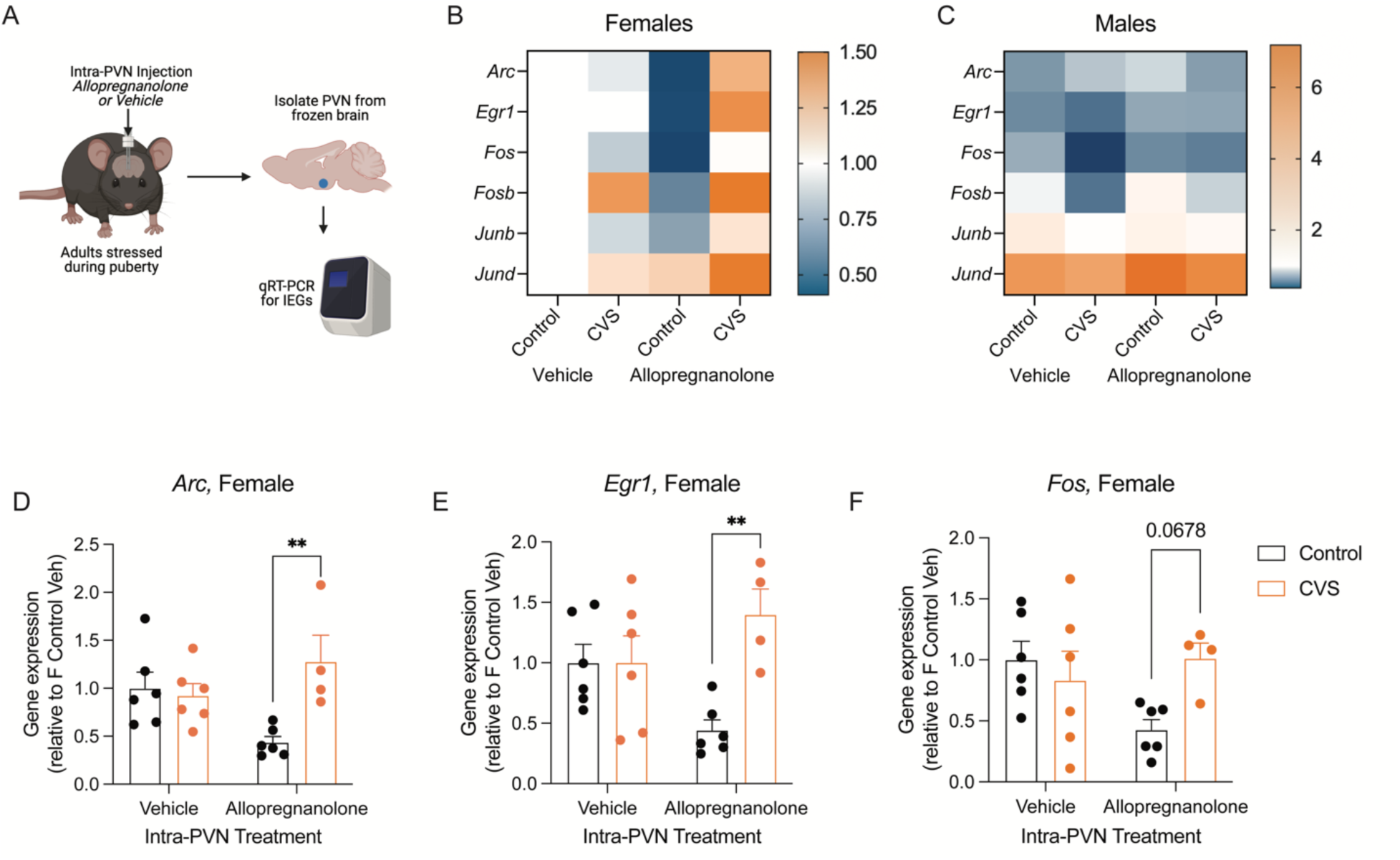
Intra-PVN allopregnanolone interacted with pubertal stress and sex to alter baseline immediate early gene expression. **(A)** Adult female and male mice that had been exposed to pubertal stress or not were cannulated so that vehicle or allopregnanolone could be microinjected into the PVN. Two hours after microinjection, brains were collected in non-stimulated conditions (baseline). The PVN was isolated, RNA was isolated, and qRT-PCR was performed for six puberty-stress associated IEGs (*Arc, Egr1, Fos, Fosb, Junb, Jund*). Created with BioRender.com. **(B)** Heat map showing expression relative to Control Vehicle females within each gene. Intra-PVN allopregnanolone generally led to a decrease in IEG expression in Control females. In contrast, CVS females responded with an increase in IEG expression. **(C)** In contrast to females, males generally had less expression of all IEGs, except for *Jund*. IEG expression in males was less sensitive to either pubertal stress or intra-PVN Allopregnanolone. **(D-F)** When IEGs were examined within sex, there was a significant interaction between pubertal stress (CVS) and intra-PVN allopregnanolone for *Arc*, *Egr1*, and *Fos*. In allopregnanolone-treated females, there was significantly increased baseline *Arc* and *Egr1* expression and a similar trend in *Fos* expression. This expression pattern recapitulates previous findings in pregnant females, where pubertally stressed females with high allopregnanolone levels in the PVN had increased IEG expression at baseline. CVS = chronic variable stress, IEG = immediate early gene, PVN = paraventricular nucleus of the hypothalamus. \*\**p* < 0.05 on Bonferroni post-hoc test. Data are mean ± SEM.

Adult female and male mice that had either been exposed to CVS or not received an intra-PVN injection of allopregnanolone or vehicle. In five of the six IEGs (*Arc, Egr1, Fos, Junb, Jund*), there was a significant effect of sex on baseline gene expression (**Table S3**). There was no effect of sex, stress, or allopregnanolone on the expression of *Fosb* (**Figure S1**). Our findings in vehicle-treated mice are consistent with our previous work, such that there were no significant differences between Control and CVS vehicle-treated adult mice (**Figure 2B,C**). The heatmaps show that males generally have decreased expression relative to vehicle treated female Controls, and the expression in males is less dynamic than in females in response to pubertal stress or intra-PVN allopregnanolone. This is consistent with our previous findings showing that males are less vulnerable to the effects of pubertal stress in the PVN and on the HPA axis than females.

Because there was an effect of sex on IEG mRNA expression, we examined the relationship between pubertal stress and intra-PVN allopregnanolone within sex. In males, there were no significant effects of pubertal stress or intra-PVN allopregnanolone on any of the IEGs (*p* > 0.05, **Table S4**). However, in females, there was a significant interaction between pubertal stress and intra-PVN allopregnanolone on baseline gene expression for *Arc* (**Figure 2D**, F(1, 18) = 8.68, *p* = 0.008), *Egr1* (**Figure 2E**, F(1, 18) = 7.54, *p* = 0.01), and *Fos* (**Figure 2F**, F(1, 18) = 4.84, *p* = 0.04). Post-hoc testing revealed a similar pattern across these three IEGs. There was no difference in baseline gene expression in vehicle-treated animals (Control vs CVS: *Arc p* > 0.9, *Egr1 p* > 0.9, *Fos p* = 0.95). Under intra-PVN allopregnanolone treatment, we replicated our prior findings from pregnant females, such that pubertally stressed females had significantly higher baseline gene expression compared to controls for *Arc* (*p* = 0.004) and *Egr1* (*p* = 0.003). There was a similar trend for *Fos* expression following allopregnanolone treatment, although this did not pass multiple comparison corrections (*p* = 0.06). The same, albeit non-significant pattern was observed in the expression of *Fosb, Junb,* and *Jund* (**Figure S1**). Thus, acute delivery of allopregnanolone into the PVN of adult females recapitulates the phenotype that we have previously observed in the brain of late pregnant females, such that pubertally stressed females have increased baseline IEG mRNA expression relative to control females. These findings are specific to females, as adult males show no significant effect of pubertal stress or intra-PVN allopregnanolone on baseline IEG mRNA expression.

### Baseline IEG mRNA expression during development is disrupted by chronic pubertal stress

To further understand the developmental origins of the baseline IEG mRNA expression patterns in pubertally stressed mice, we measured baseline gene expression in the PVN at three time points during the age range when CVS is applied (PN21, PN28, PN35, **Figure 3A**). We first examined whether there were any sex differences in baseline gene expression at PN21, prior to the onset of pubertal stress. At PN21, there were no differences between females and males in baseline gene expression (all t-test *p* > 0.05, **Table S5**). We then examined whether each gene had any variability by sex in baseline expression in Control animals at the three ages examined. Linear regression analysis showed that there was limited impact of sex on baseline mRNA expression from PN21 to PN35 in Controls (**Figure 3B, Table S6**). Only in *Arc* was there a significant change in gene expression, and only in males (F (1,24) = 10.29, *p* = 0.003), whose baseline *Arc* expression increased with age. Otherwise, linear regressions showed that both female and male Controls had stable baseline IEG mRNA expression from PN21 to PN35 (*p* > 0.05).

**Figure 3.**
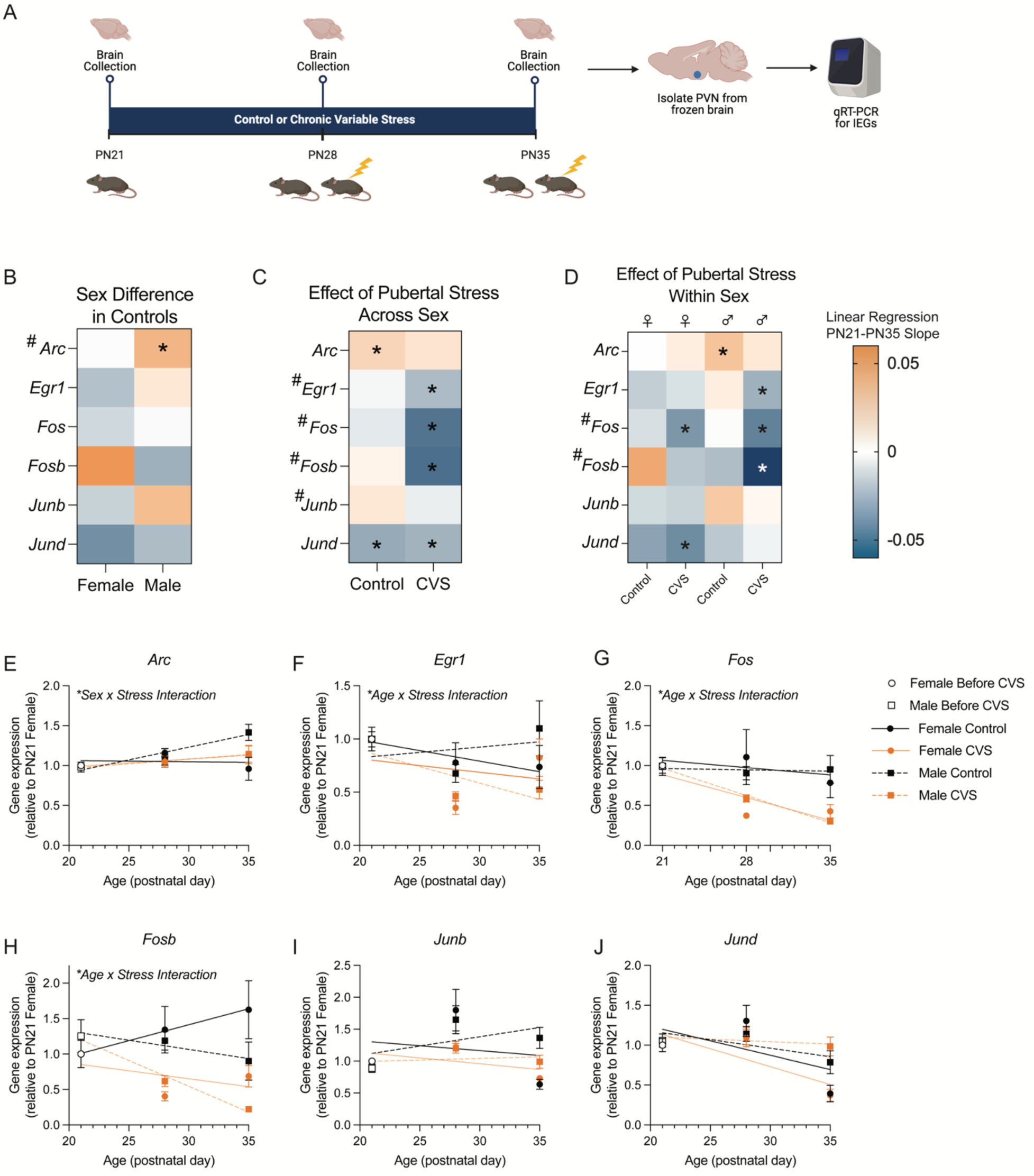
Baseline immediate early gene expression was dynamic during early adolescence and was altered by pubertal stress. **(A)** Brains were collected from early adolescent female and male mice that were exposed to pubertal stress or not at three time points. These time points represent prior to starting CVS (PN21), after 1 week of CVS (PN28), and after two weeks of CVS (PN35). Brains were collected 24 hours after the prior stressor and in non-stimulated conditions. The PVN was dissected, RNA was isolated, and qRT-PCR was performed for six puberty-stress associated immediate early genes (IEG; *Arc, Egr1, Fos, Fosb, Junb, Jund*). Created with BioRender.com. **(B-D)** Asterisks (*) indicate a slope of the regression line from PN21-Pn35 is significantly non-zero, while pound signs (#) indicate that the slopes are significantly different between the groups within a gene. **(B)** Heat map depicting the direction of the slope of the linear regression line calculated from PN21 to PN35 for gene expression in the PVN of Control (female and male) mice. The heat map shows that there were few sex differences in the trajectory of expression in control mice during this window, with only *Arc* showing a difference between sexes. **(C)** Heat map depicting direction of the slope of the linear regression line for gene expression in the PVN based on pubertal stress exposure. Here, males and females were grouped together. **(D)** Heat map depicting the direction of the slope of the linear regression line calculated from PN21 to PN35 for gene expression in the PVN of Control (female and male) and CVS (female and male) mice. The heat map shows that CVS disrupts the trajectories of each of the six IEGs. **(C)** Heat map that depicts the direction of the slope on the linear regression line based on pubertal stress and sex. There was a sex difference in the impact of pubertal stress on the trajectory of baseline IEG expression during adolescence for several genes. **(E-J)** Gene expression and linear regression lines for all six genes at PN21, PN28, and PN35. Significant interactions between factors (sex, pubertal stress, age) are indicated where appropriate. CVS = chronic variable stress, IEG = immediate early gene, PN = postnatal day, PVN = paraventricular nucleus of the hypothalamus. Data are mean ± SEM.

Next, we examined the impact of pubertal stress using linear regression analyses. When sexes were collapsed, we found an impact of pubertal stress on the trajectory of gene expression from PN21 to PN35 for *Egr1*, *Fos, Fosb,* and *Junb*, but not for *Arc* or *Jund* (**Figure 3C, Figure S2, Table S7**). In general, pubertal stress decreased baseline IEG mRNA expression for those genes affected. When we disaggregated sex to examine the impact of pubertal stress in a sex-specific way, we found that some effects of pubertal stress were sex-specific (**Figure 3D, Tables S8-S11**).

Multiple regression analysis showed that there was an interaction between sex and stress on baseline *Arc* expression (*p* = 0.02), although this interaction was not supported in the parallel 3-way ANOVA analysis, which revealed only a trend for a sex x stress interaction (*p* = 0.06). For *Arc*, female mice had no change in baseline gene expression throughout early adolescence and pubertal stress had no effect (**Figure 3E**). Control male mice showed a slight increase in *Arc* throughout early adolescence that was absent in pubertally stressed mice.

For *Egr1* (**Figure 3F**), there was a significant interaction between age, sex, and stress on baseline expression (F(1,66) = 4.27, *p* = 0.04). In Controls, females tended to have a decrease in *Egr1* expression throughout early adolescence, whereas males had an increase. In both sexes, exposure to pubertal stress decreased baseline *Egr1* expression throughout the early adolescent window.

For *Fos* (**Figure 3G**), *Fosb* (**Figure 3H**), and *Junb* (**Figure 3I**), there was a main effect of pubertal stress, such that pubertal stress led to a decrease in gene expression in both sexes (*Fos*: F(1,61) = 20.71, *p* < 0.0001; *Fosb*: F(1,61) = 31.37, *p* < 0.0001; *Junb*: F(1,64) = 7.15, *p* = 0.0009). Multiple regression analysis pointed to an interaction between stress and age for *Fos* (*p* < 0.0001) and *Fosb* (*p* < 0.0001), which shows that the magnitude of the decrease in expression in CVS mice was larger at PN35 than at PN28.

Finally, there were no effects of pubertal stress on *Jund* expression from PN21 through PN35 (**Figure 3J**). There was an interaction between age and sex (F(1,66) = 9.63, *p* = 0.003) on baseline expression of *Jund*, such that the difference between expression in males and females became larger at PN35 than it was at PN28, when there were no differences between sexes.

Overall, these analyses show that baseline IEG mRNA expression was influenced in an interactive way by age, stress, and pubertal stress, although not always identically across IEGs.

## Discussion

Why individuals have negative consequences following stress is a complex phenomenon that is dictated by individual factors, when in the lifespan the stress is experienced, and when the consequences are measured, both proximity to the stressful event and timepoint within the lifespan (30–35). For women, who are more likely than men to suffer from affective disorders, risk factors include experiencing stressful or traumatic events around the onset of puberty and experiencing later times of hormonal change such as pregnancy and aging (36,37). We have previously shown that in both humans and mice, adversity experienced around the onset of puberty led to a blunted hypothalamic-pituitary-adrenal (HPA) stress response during the peripartum window and increased reporting of postpartum depressive symptoms in humans (9). We have used this translationally-relevant mouse model to investigate the mechanisms underlying the lasting effect of pubertal stress on the HPA axis response during the peripartum window. Our previous work identified latent epigenetic changes in the paraventricular nucleus of the hypothalamus (PVN) that are uncovered by the experience of pregnancy, potentially by the increased allopregnanolone levels (11). Here, we expanded on those findings to directly link allopregnanolone action in the PVN to the molecular changes associated with pubertal stress.

Our previous analysis of the PVN transcriptome revealed increases in the baseline gene expression of six immediately early genes (IEGs) in pregnant females as a result of pubertal adversity (9). We utilized PVN-specific pharmacological manipulations to address whether acute allopregnanolone was sufficient to induce an increase in baseline IEG mRNA expression in pubertally stressed mice. In adult female mice, a single dose of allopregnanolone administered into the PVN recapitulated the increased baseline IEG expression only in pubertally stressed females. Allopregnanolone is a neurosteroid metabolite of progesterone that is present at high levels in the brain during pregnancy. It binds to several receptors, including extrasynaptic GABA-A, pregnane X receptors, and progesterone receptors, although its function as an FDA approved treatment for postpartum depression is attributed to its binding to the GABA-A receptor (16–18,38). In our studies, allopregnanolone does not act to reduce risk for negative outcomes in prenatal mice. We previously showed that allopregnanolone is both necessary and sufficient to uncover the blunted HPA axis response, suggesting that it is an agent of risk during late pregnancy in mice with a history of stress during puberty (11). It is possible that the effect of allopregnanolone in altering the transcriptome in response to pubertal stress is working via a GABA system that has been altered by pubertal stress or via different mechanisms, including the transcription-regulating pregnane X receptor (39).

One thing that both puberty and pregnancy have in common is a dynamic change in allopregnanolone levels. There is an increase in allopregnanolone during puberty in both humans and rodents that has been suggested to be important for the normative development of the HPA axis (40,41) and which parallels both the increase and regulatory role of allopregnanolone in pregnancy. Clinical studies have shown allopregnanolone levels during the second trimester are predictive of postpartum depression and anxiety symptoms (42,43). Our work suggests that pubertal stress alters the transcriptional and physiological response to allopregnanolone levels. Future work will need to resolve whether individuals who have a history of preconception adversity are likely to fully benefit from neurosteroid-based antidepressants.

These findings provide further evidence that increased baseline IEG mRNA expression may be a key molecular switch that is programmed by pubertal stress and leads to the vulnerability of an altered HPA axis in adulthood, during which adult changes in allopregnanolone can impinge upon IEG mRNA expression to uncover the vulnerability. However, whether this effect is programmed immediately after pubertal stress, at postnatal day 35, or whether it develops in the weeks between PN35 and adulthood, was unknown. Understanding the developmental origins of this molecular phenotype could lead to early interventions and preventative approaches for negative consequences typically identified in adulthood (44). Therefore, we examined baseline IEG mRNA expression in the PVN during the window of pubertal stress in females and males. First, we examined developmental changes in gene expression in female and male mice. Overall, baseline IEG mRNA expression in the PVN was relatively stable from PN21-PN35 in controls. Males and females did not differ in gene expression at PN21, but often showed different trajectories of gene expression when examined 1 week (PN28) or 2 weeks (PN35) later. These data show that baseline IEG mRNA expression in the PVN is not static. We further found that pubertal stress altered IEG expression during this adolescent window. In general, the effect of pubertal stress was to decrease baseline IEG mRNA expression in both females and males.

Future studies will resolve these findings in a cell-type specific manner. In our current and past studies, we have examined the complete PVN. While limited work has addressed cell-type specific outcomes in the hypothalamus following adolescent stress, there is work from early prenatal stress that informs future cell-type specific studies. For example, limited bedding and nesting produced alterations to the transcriptome of CRF cells in 1.5 week old mice, such that CRF cells that co-express glutamate, but not GABA, may have enhanced sensitivity to this early life adversity (45). This is consistent with prior work showing increased synaptic glutamatergic drive onto neonatal PVN CRF neurons following limited nesting (46). This study also showed that mice exposed to limited nesting had altered sensitivity to allopregnanolone. It is possible that what we and others are finding is a mechanism wherein CRF cells are being tuned by both glutamatergic and GABAergic systems, and that early life and adolescent stressors impinge upon both neurotransmitter systems to lead to vulnerability to negative outcomes later in life.

Another future goal will be to merge this cell-type specificity with tracking of the same cells throughout the lifespan, which will overcome our current limitation of examining adolescent and adult outcomes in different animals. Recent work has shown that a stress exposure early, in the form of maternal separation, led to the prolonged sensitivity of a set of cells in the nucleus accumbens to stressors in adulthood (47). Balouek et al showed that neurons transgenically tagged by maternal separation were the same cells reactivated by stress later in life and that blocking their activation ameliorated the negative behavioral consequences. Interestingly, it was males that were sensitive to the adult manipulations. This contrasts with our study and others conducted in adolescence, where females tend to be more vulnerable to lasting negative outcomes of stress exposure.

While our model considers times in the lifespan when females are more vulnerable than males, we included male mice in all experiments here. Consistent with our previous work showing that adult females are more vulnerable to the lasting effects of pubertal stress, we found that baseline IEG mRNA expression in males is less sensitive in adulthood than it is in females. This contrasts with our finding that adolescent males display changes in baseline IEG mRNA expression the PVN in response to pubertal stress during the stress window. Future studies will resolve why the molecular responsiveness in the PVN of males does not also result in negative outcomes in adulthood and will consider other brain regions and behavioral outcomes that might provide evidence of negative plasticity in males. Altogether, our study and others conducted recently continue to lend support for to the literature on sex differences in vulnerability to early life stress (48–51). Ongoing studies will continue to refine our knowledge on the molecular underpinnings of how sex and early life stress interact to produce vulnerability to negative outcomes.

Few studies have examined the expression levels of IEGs in non-stimulated (baseline) conditions either after acute or repeated stimulus presentation. Studies where baseline IEG mRNA or protein have been measured have been focused on the impact of neonatal experiences. Embryonic rearing conditions of chick eggs influences lateralization of spontaneous (baseline) c-Fos protein expression in several brain regions (52). Another study that examined the regulators of baseline *Arc* mRNA expression showed that blocking brain-derived neurotrophic factor reduced baseline *Arc* expression in the neocortex, which gives insight into molecules that may participate in the lasting effects that we have observed here (53). Repeatedly evoking seizures in rats results in blunted IEG mRNA expression in the hippocampus in response to a locomotor task during a seizure-free period (54), although differences in protein levels were not detectable in home-cage controls (55). In our model, four of the six IEGs are from the Jun and Fos family of proteins, which dimerize to form the AP-1 complex in order to regulate transcription (56). Binding of the AP-1 complex is required in the recruitment of the glucocorticoid receptor (57). It is possible that the lasting HPA axis physiological plasticity observed in pregnant, pubertally stressed females is driven by alterations to the AP-1 complex that subsequently impact glucocorticoid signaling. Alternatively, it is possible that altered baseline IEG expression influences the cellular response of CRF cells to PVN inputs that initiate the HPA axis response. Future studies will address where in the cascade the IEGs may be influencing that glucocorticoid response to stress.

There are competing hypotheses for the way in which multiple stressors throughout the lifespan interact to influence risk for negative outcomes. Some findings support a mismatch theory, where organisms that experience early life stress are well suited to future stressful environments and appear resilient (58). Our prior results and new findings support the idea of cumulative stress, such that a ‘second hit’ challenge results in an increased risk for negative outcomes, also termed an increased allostatic load (59). However, when humans and animals experience that ‘second hit’ of challenge, whether that is another exposure to stress later in life or to physiological changes such as pregnancy, it has not been clear what the molecular agents of that ‘hit’ are. We have identified that allopregnanolone is a molecular agent of that ‘second hit’ of risk and that it interacts with IEGs to influence their baseline gene expression (Figure 4). This interaction at baseline could lead to differences in how the cells in the PVN respond to stimuli and could underlie the blunted HPA axis response and altered behavior we have previously observed.

**Figure 4.**
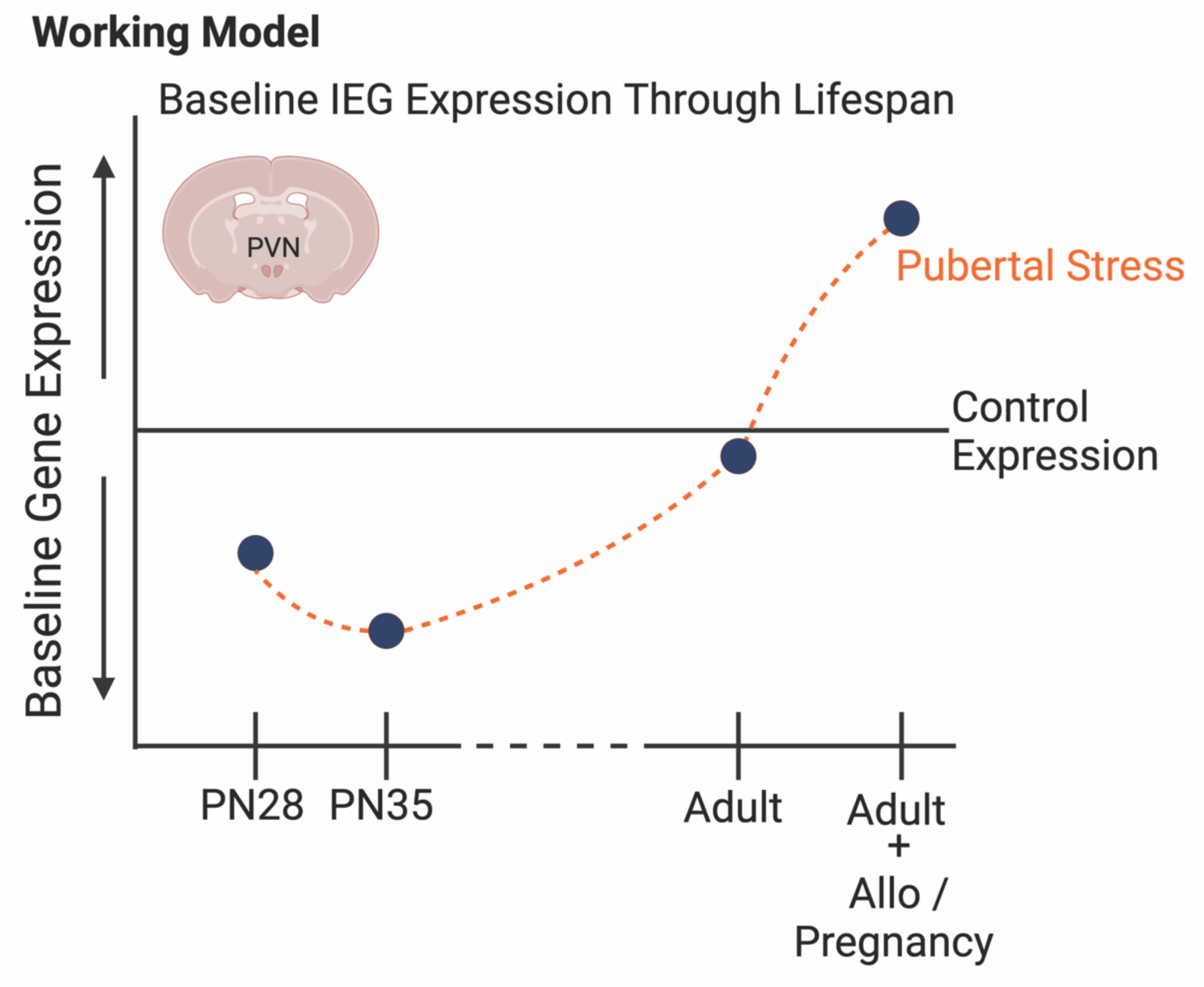
Schematic summarizing our findings on the influence of pubertal stress on the transcriptome of the developing and adult hypothalamus. Through prior and current findings, we have demonstrated that baseline gene expression for a set of immediate early genes in the paraventricular nucleus of the hypothalamus is sensitive to pubertal stress (chronic variable stress from postnatal days 21-34) both during the stress exposure and in adulthood. Of note, all IEG mRNA expression differences were measured in non-stimulated, baseline conditions, and so may represent how the PVN is poised to respond to stimuli. Generally, baseline IEG mRNA expression is decreased during the pubertal stress time period in CVS mice relative to Controls. Whether adult mice are pregnant or are exposed to allopregnanolone, there is an increase in baseline IEG expression in females. This suggests that the PVN is poised to respond differently, which we have observed in the form of a blunted glucocorticoid response to restraint in adulthood under allopregnanolone or pregnancy. Our new findings provide support for a role of baseline IEG expression in the mechanism underlying the lasting effect of pubertal stress on the PVN. Created with BioRender.com.

We have taken a developmental origins approach to the question of how stress during puberty impacts an individual through the lifespan, wherein we are generating a trajectory of molecular, physiological, and behavioral outcomes both during early life stress and into adulthood. Our findings show that a key molecular outcome associated with the negative adult outcomes after pubertal stress – the permissive expression of a set of IEG mRNA at baseline - is plastic, displaying normative changes throughout development, and that stress during puberty results in a disruption of baseline expression that is detectable as early as adolescence. These studies provide novel insight into the potential mechanisms underlying female-relevant risk for stress dysregulation, a central endophenotype of affective disorders. Understanding the early signs and mechanisms of negative outcomes like stress dysregulation and affective dysfunction that, both in humans and animal models, are generally identified and treated in adulthood, will lead to early interventions, preventative approaches, and better outcomes for lifetime health.

## Author Contributions

KEM designed experiments. KNG, SLH, JMM, and KEM conducted experiments, sorted, and analyzed data. KNG wrote an initial draft of the manuscript, which was edited to a final version by KEM.

## Acknowledgments

This work was supported by National Institutes of Health Grant R00 HD091376 (KEM) and start-up funds to KEM. KNG and JMM were additionally funded by the West Virginia University Summer Undergraduate Research Experience program.

## Data Availability

Data will be publicly available via the Open Science Framework repository upon publication.

