## Supplemental Material for "The impact of pubertal stress and adult hormone exposure on the transcriptome of the developing hypothalamus"

#### Supplementary Material

### Supplemental Tables

Table S1. Final sample sizes by group

| Sex | Pubertal Stress | Intra-PVN Tx | Starting Group Size | Final Group Size | Final Number of Litters | Reason for Attrition |
|---|---|---|---|---|---|---|
| **Experiment 1** | | | | | | |
| Female | Control | Vehicle | 7 | 6 | 5 | headcap complication (n=1) |
| Female | CVS | Vehicle | 7 | 6 | 5 | RNA quality (n=1) |
| Female | Control | Allopregnanolone | 7 | 7 | 6 | n/a |
| Female | CVS | Allopregnanolone | 5 | 4 | 3 | misplaced cannula (n=1) |
| Male | Control | Vehicle | 5 | 5 | 4 | n/a |
| Male | CVS | Vehicle | 4 | 4 | 3 | n/a |
| Male | Control | Allopregnanolone | 4 | 4 | 4 | n/a |
| Male | CVS | Allopregnanolone | 6 | 5 | 4 | misplaced cannula (n=1) |
| **Experiment 2** | | | | | | |
| **Sex** | **Pubertal Stress** | **Age of Collection** | **Starting Group Size** | **Final Group Size** | **Final Number of Litters** | **Reason for Attrition** |
| Female | Control | PN21 | 10 | 9 | 9 | sample collection issue (n=1) |
| Female | Control | PN28 | 10 | 10 | 8 | n/a |
| Female | CVS | PN28 | 10 | 10 | 10 | n/a |
| Female | Control | PN35 | 10 | 6 | 6 | RNA isolation error (n=4) |
| Female | CVS | PN35 | 10 | 9 | 9 | cryostat issue (n=1) |
| Male | Control | PN21 | 9 | 7 | 7 | sample collection issue (n=2) |
| Male | Control | PN28 | 10 | 10 | 10 | n/a |
| Male | CVS | PN28 | 10 | 10 | 10 | n/a |
| Male | Control | PN35 | 10 | 10 | 9 | n/a |
| Male | CVS | PN35 | 10 | 10 | 10 | n/a |

*Tx = treatment via microinjection; CVS = chronic variable stress; PN = postnatal day

Table S2. qRT-PCR gene information

| Gene | Gene Symbol | TaqMan Assay ID | RefSeq Reference Sequence |
|---|---|---|---|
| Activity regulated cytoskeletal-associated protein | *Arc* | Mm01204954_g1 | NM_001276684.1 |
| Early growth response 1 | *Egr1* | Mm00656724_m1 | NM_007913.5 |
| FBJ osteosarcoma oncogene | *Fos* | Mm00487425_m1 | NM_010234.2 |
| FBJ osteosarcoma oncogene B | *Fosb* | Mm00500401_m1 | NM_008036.2 |
| Glyceraldehyde-3-phosphate dehydrogenase | *Gapdh* | Mm99999915_g1 | NM_001289726.1 |
| Jun B proto-oncogene | *Junb* | Mm04243546_s1 | NM_008416.3 |
| Jun D proto-oncogene | *Jund* | Mm00495088_s1 | NM_001286944.1 |

Table S3. Full statistical reporting from 3-way ANOVA analysis in Experiment 1

| Gene | Factor | F value | P value | Significance |
|---|---|---|---|---|
| *Arc* | | | | |
| | Drug | F (1, 32) = 0.08 | P=0.78 | ns |
| | Sex | F (1, 32) = 2.12 | P=0.15 | ns |
| | Stress | F (1, 32) = 2.64 | P=0.11 | ns |
| | Drug x Sex | F (1, 32) = 0.42 | P=0.52 | ns |
| | Drug x Stress | F (1, 32) = 1.64 | P=0.21 | ns |
| | Sex x Stress | F (1, 32) = 3.17 | P=0.08 | ns |
| | **Drug x Sex x Stress** | F (1, 32) = 7.85 | P=0.008 | * |
| *Egr1* | | | | |
| | Drug | F (1, 32) = 0.05 | P=0.82 | ns |
| | **Sex** | F (1, 32) = 7.97 | P=0.008 | * |
| | Stress | F (1, 32) = 3.51 | P=0.07 | ns |
| | Drug x Sex | F (1, 32) = 0.83 | P=0.37 | ns |
| | **Drug x Stress** | F (1, 32) = 4.91 | P=0.03 | * |
| | **Sex x Stress** | F (1, 32) = 5.10 | P=0.03 | * |
| | Drug x Sex x Stress | F (1, 32) = 3.60 | P=0.07 | ns |
| *Fos* | | | | |
| | Drug | F (1, 32) = 0.49 | P=0.49 | ns |
| | **Sex** | F (1, 32) = 5.00 | P=0.03 | * |
| | Stress | F (1, 32) = 0.007 | P=0.93 | ns |
| | Drug x Sex | F (1, 32) = 1.14 | P=0.29 | ns |
| | **Drug x Stress** | F (1, 32) = 5.66 | P=0.02 | * |
| | Sex x Stress | F (1, 32) = 3.22 | P=0.08 | ns |
| | Drug x Sex x Stress | F (1, 32) = 0.10 | P=0.32 | ns |
| *Fosb* | | | | |
| | Drug | F (1, 31) = 0.24 | P=0.63 | ns |
| | Sex | F (1, 31) = 0.28 | P=0.60 | ns |
| | Stress | F (1, 31) = 0.02 | P=0.90 | ns |
| | Drug x Sex | F (1, 31) = 0.97 | P=0.33 | ns |
| | Drug x Stress | F (1, 31) = 0.04 | P=0.84 | ns |
| | Sex x Stress | F (1, 31) = 3.87 | P=0.06 | ns |
| | Drug x Sex x Stress | F (1, 31) = 0.40 | P=0.53 | ns |
| *Junb* | | | | |
| | Drug | F (1, 32) = 0.03 | P=0.87 | ns |
| | **Sex** | F (1, 32) = 10.09 | P=0.003 | * |
| | Stress | F (1, 32) = 1.15 | P=0.29 | ns |
| | Drug x Sex | F (1, 32) = 0.002 | P=0.96 | ns |
| | Drug x Stress | F (1, 32) = 2.12 | P=0.15 | ns |
| | Sex x Stress | F (1, 32) = 3.19 | P=0.08 | ns |
| | Drug x Sex x Stress | F (1, 32) = 0.001 | P=0.97 | ns |
| *Jund* | | | | |
| | Drug | F (1, 32) = 2.18 | P=0.15 | ns |
| | **Sex** | F (1, 32) = 77.90 | P<0.0001 | * |
| | Stress | F (1, 32) = 0.27 | P=0.60 | ns |
| | Drug x Sex | F (1, 32) = 1.00 | P=0.32 | ns |
| | Drug x Stress | F (1, 32) = 0.01 | P=0.91 | ns |
| | Sex x Stress | F (1, 32) = 0.81 | P=0.38 | ns |
| | Drug x Sex x Stress | F (1, 32) = 0.087 | P=0.77 | ns |

*Outcome: gene expression relative to Control Vehicle Female; ns = not significant; *p < 0.05; Drug (Vehicle, Allopregnanolone); Sex (Female, Male); Stress (Control, CVS)*

Table S4. Full statistical reporting from within-sex 2-way ANOVA analysis in Experiment 1

| Gene | Factor | F value | P value | Significance |
|---|---|---|---|---|
| *Arc* | | | | |
| Female | Drug | F (1, 18) = 0.46 | P=0.51 | ns |
| | **Stress** | F (1, 18) = 6.05 | P=0.02 | * |
| | **Drug x Stress** | F (1, 18) = 8.68 | P=0.009 | * |
| Male | Drug | F (1, 14) = 0.06 | P=0.80 | ns |
| | Stress | F (1, 14) = 0.01 | P=0.91 | ns |
| | Drug x Stress | F (1, 14) = 1.14 | P=0.30 | ns |
| *Egr1* | | | | |
| Female | Drug | F (1, 18) = 0.21 | P=0.65 | ns |
| | **Stress** | F (1, 18) = 7.60 | P=0.01 | * |
| | **Drug x Stress** | F (1, 18) = 7.54 | P=0.01 | * |
| Male | Drug | F (1, 14) = 0.85 | P=0.37 | ns |
| | Stress | F (1, 14) = 0.10 | P=0.76 | ns |
| | Drug x Stress | F (1, 14) = 0.07 | P=0.80 | ns |
| *Fos* | | | | |
| Female | Drug | F (1, 18) = 1.33 | P=0.26 | ns |
| | Stress | F (1, 18) = 1.49 | P=0.24 | ns |
| | **Drug x Stress** | F (1, 18) = 4.84 | P=0.04 | * |
| Male | Drug | F (1, 14) = 0.10 | P=0.76 | ns |
| | Stress | F (1, 14) = 2.17 | P=0.16 | ns |
| | Drug x Stress | F (1, 14) = 1.41 | P=0.25 | ns |
| *Fosb* | | | | |
| Female | Drug | F (1, 17) = 0.17 | P=0.68 | ns |
| | Stress | F (1, 17) = 3.16 | P=0.09 | ns |
| | Drug x Stress | F (1, 17) = 0.50 | P=0.49 | ns |
| Male | Drug | F (1, 14) = 0.79 | P=0.39 | ns |
| | Stress | F (1, 14) = 1.22 | P=0.29 | ns |
| | Drug x Stress | F (1, 14) = 0.07 | P=0.80 | ns |
| *Junb* | | | | |
| Female | Drug | F (1, 18) = 0.05 | P=0.83 | ns |
| | Stress | F (1, 18) = 0.52 | P=0.48 | ns |
| | Drug x Stress | F (1, 18) = 2.02 | P=0.17 | ns |
| Male | Drug | F (1, 14) = 0.004 | P=0.95 | ns |
| | Stress | F (1, 14) = 2.40 | P=0.14 | ns |
| | Drug x Stress | F (1, 14) = 0.66 | P=0.43 | ns |
| *Jund* | | | | |
| Female | Drug | F (1, 18) = 1.70 | P=0.21 | ns |
| | Stress | F (1, 18) = 1.05 | P=0.32 | ns |
| | Drug x Stress | F (1, 18) = 0.23 | P=0.63 | ns |
| Male | Drug | F (1, 14) = 1.33 | P=0.27 | ns |
| | Stress | F (1, 14) = 0.44 | P=0.52 | ns |
| | Drug x Stress | F (1, 14) = 0.04 | P=0.85 | ns |

*Outcome: gene expression relative to Control Vehicle Female; ns = not significant; *p < 0.05; Drug (Vehicle, Allopregnanolone); Sex (Female, Male); Stress (Control, CVS)*

Table S5. Results of t-testing for analysis of sex effect on gene expression at PN21 in Experiment 2

| Gene | t ratio | df | P value | Significance |
|---|---|---|---|---|
| *Arc* | 0.32 | 13.00 | 0.75 | ns |
| *Egr1* | 0.07 | 14.00 | 0.95 | ns |
| *Fos* | 0.09 | 14.00 | 0.93 | ns |
| *Fosb* | 0.55 | 13.00 | 0.59 | ns |
| *Junb* | 1.38 | 14.00 | 0.19 | ns |
| *Jund* | 0.48 | 14.00 | 0.63 | ns |

*Outcome: gene expression relative to PN21 Females; ns = not significant*

Table S6. Results of linear regression in Controls for analysis of sex differences in slopes of gene expression from PN21-PN35 in Experiment 2

| Gene | Factor | F | P value | Significance |
|---|---|---|---|---|
| *Arc* | | | | |
| | Female slope | F (1,21) = 0.03 | 0.86 | ns |
| | **Male slope** | F (1,24) = 10.29 | 0.004 | * |
| | **Female vs Male** | F (1,45) = 6.24 | 0.02 | * |
| *Egr1* | | | | |
| | Female slope | F (1,23) = 1.23 | 0.28 | ns |
| | Male slope | F (1,24) = 0.28 | 0.60 | ns |
| | Female vs Male | F (1,47) = 1.31 | 0.26 | ns |
| *Fos* | | | | |
| | Female slope | F (1,22) = 0.25 | 0.62 | ns |
| | Male slope | F (1,23) = 0.03 | 0.86 | ns |
| | Female vs Male | F (1,45) = 0.14 | 0.71 | ns |
| *Fosb* | | | | |
| | Female slope | F (1,21) = 2.21 | 0.15 | ns |
| | Male slope | F (1,23) = 1.36 | 0.25 | ns |
| | Female vs Male | F (1,44) = 3.65 | 0.06 | ns |
| *Junb* | | | | |
| | Female slope | F (1,22) = 0.27 | 0.60 | ns |
| | Male slope | F (1,25) = 1.99 | 0.17 | ns |
| | Female vs Male | F (1,47) = 1.59 | 0.21 | ns |
| *Jund* | | | | |
| | Female slope | F (1,23) = 3.34 | 0.08 | ns |
| | Male slope | F (1,25) = 2.87 | 0.10 | ns |
| | Female vs Male | F (1,48) = 0.40 | 0.53 | ns |

*Outcome: gene expression relative to PN21 Females; ns = not significant*

Table S7. Results of linear regression for analysis of impact of pubertal stress on slopes of gene expression from PN21-PN35 in Experiment 2

| Gene | Factor | F | P value | Significance |
|---|---|---|---|---|
| *Arc* | | | | |
| | **Control slope** | F (1,47) = 6.58 | 0.01 | * |
| | CVS slope | F (1,52) = 3.31 | 0.07 | ns |
| | Control vs CVS slope | F (1,99) = 0.63 | 0.43 | ns |
| | Control vs CVS intercept | F (1,100) = 1.69 | 0.20 | ns |
| *Egr1* | | | | |
| | Control slope | F (1,49) = 0.08 | 0.78 | ns |
| | **CVS slope** | F (1,53) = 5.90 | 0.02 | * |
| | Control vs CVS slope | F (1,102) = 1.40 | 0.24 | ns |
| | **Control vs CVS intercept** | F (1,103) = 5.68 | 0.02 | * |
| *Fos* | | | | |
| | Control slope | F (1,47) = 0.31 | 0.58 | ns |
| | **CVS slope** | F (1,50) = 59.83 | < 0.0001 | * |
| | **Control vs CVS slope** | F (1,97) = 6.21 | 0.01 | * |
| *Fosb* | | | | |
| | Control slope | F (1,46) = 0.097 | 0.76 | ns |
| | **CVS slope** | F (1,51) = 18.57 | < 0.0001 | * |
| | **Control vs CVS slope** | F (1,97) = 6.02 | 0.02 | * |
| *Junb* | | | | |
| | Control slope | F (1,49) = 0.33 | 0.57 | ns |
| | CVS slope | F (1,51) = 0.85 | 0.36 | ns |
| | Control vs CVS slope | F (1,100) = 0.80 | 0.37 | ns |
| | **Control vs CVS intercept** | F (1,101) = 6.74 | 0.01 | * |
| *Jund* | | | | |
| | **Control slope** | F (1,50) = 6.04 | 0.02 | * |
| | **CVS slope** | F (1,52) = 7.83 | 0.007 | * |
| | Control vs CVS slope | F (1,102) = 0.04 | 0.84 | ns |
| | Control vs CVS intercept | F (1,103) = 0.22 | 0.64 | ns |

*Outcome: gene expression collapsed across sexes (both male and Female collapsed) relative to Control PN21; ns = not significant; *p < 0.05; CVS = Chronic Variable Stress*

Table S8. Results of linear regression for analysis of sex-specific impact of pubertal stress on slopes of gene expression from PN21-PN35 in Experiment 2

| Gene | Factor | F | P value | Significance |
|---|---|---|---|---|
| *Arc* | | | | |
| | Female Control | F (1,21) = 0.03 | 0.86 | ns |
| | Female CVS | F (1,25) = 1.34 | 0.26 | ns |
| | **Male Control** | F (1,24) = 10.29 | 0.004 | * |
| | Male CVS | F (1, 25) = 1.87 | 0.18 | ns |
| | Slope difference | F (3,95) = 2.33 | 0.08 | ns |
| | Intercept difference | F (3,98) = 1.48 | 0.22 | ns |
| *Egr1* | | | | |
| | Female Control | F (1,23) = 1.23 | 0.28 | ns |
| | Female CVS | F (1,26) = 0.66 | 0.42 | ns |
| | Male Control | F (1,24) = 0.28 | 0.60 | ns |
| | **Male CVS** | F (1,25) = 12.02 | 0.002 | * |
| | Slope difference | F (3,98) = 1.21 | 0.31 | ns |
| | Intercept difference | F (3,101) = 2.15 | 0.10 | ns |
| *Fos* | | | | |
| | Female Control | F (1,22) = 0.25 | 0.62 | ns |
| | **Female CVS** | F (1,25) = 19.05 | 0.0002 | * |
| | Male Control | F (1,23) = 0.03 | 0.86 | ns |
| | **Male CVS** | F (1,23) = 54.38 | <0.0001 | * |
| | Slope difference | F (3,93) = 2.15 | 0.09 | ns |
| | **Intercept difference** | F (3,96) = 5.73 | 0.001 | * |
| *Fosb* | | | | |
| | Female Control | F (1,21) = 2.21 | 0.15 | ns |
| | Female CVS | F (1,25) = 1.92 | 0.18 | ns |
| | Male Control | F (1,23) = 1.36 | 0.25 | ns |
| | **Male CVS** | F (1,24) = 34.34 | <0.0001 | * |
| | **Slope difference** | F (3,93) = 5.19 | 0.002 | * |
| *Junb* | | | | |
| | Female Control | F (1,22) = 0.27 | 0.61 | ns |
| | Female CVS | F (1,25) = 3.86 | 0.06 | ns |
| | Male Control | F (1,25) = 1.99 | 0.17 | ns |
| | Male CVS | F (1,24) = 0.25 | 0.62 | ns |
| | Slope difference | F (3,96) = 1.42 | 0.24 | ns |
| | Intercept difference | F (3,99) = 2.53 | 0.06 | ns |
| *Jund* | | | | |
| | Female Control | F (1,23) = 3.34 | 0.08 | ns |
| | **Female CVS** | F (1,25) = 16.09 | 0.0005 | * |
| | Male Control | F (1,25) = 2.87 | 0.10 | ns |
| | Male CVS | F (1,25) = 0.29 | 0.60 | ns |
| | Slope difference | F (3,98) = 1.56 | 0.20 | ns |
| | Intercept difference | F (3,101) = 1.70 | 0.12 | ns |

*Outcome: gene expression collapsed across sexes (both male and female) relative to Control PN21; ns = not significant; *p < 0.05; CVS = Chronic Variable Stress*

Table S9. Regression model results of multiple linear regression for analysis of the impact of age, sex, and stress on gene expression from PN21-PN35 in Experiment 2

| Gene | Model | SS | DF | MS | F value | P value | Significance |
|---|---|---|---|---|---|---|---|
| *Arc* | | | | | | | |
| | **2-way** | 0.7526 | 3 | 0.2509 | F (3, 99) = 3.77 | P=0.01 | * |
| *Egr1* | | | | | | | |
| | **2-way** | 1.692 | 3 | 0.5639 | F (3, 102) = 2.90 | P=0.04 | * |
| *Fos* | | | | | | | |
| | **2-way** | 5.549 | 3 | 1.850 | F (3, 97) = 10.02 | P<0.0001 | * |
| *Fosb* | | | | | | | |
| | **2-way** | 10.67 | 3 | 3.556 | F (3, 97) = 10.14 | P<0.0001 | * |
| *Junb* | | | | | | | |
| | **2-way** | 2.469 | 3 | 0.8229 | F (3, 100) = 3.11 | P=0.03 | * |
| *Jund* | | | | | | | |
| | 2-way | 1.122 | 3 | 0.3741 | F (3, 101) = 2.16 | P=0.10 | ns |
| | **Main effects** | 2.632 | 3 | 0.8774 | F (3, 101) = 5.54 | P=0.001 | * |

*Outcome: gene expression relative to Control PN21Female; Analysis was used to determine the most appropriate regression model to utilize; ns = not significant; *p < 0.05*

Table S10. Multiple linear regression parameter estimates for analysis of the impact of age, sex, and stress on gene expression from PN21-PN35 in Experiment 2

| Gene | Variable | B | SE | 95% CI | t value | P value | Significance |
|---|---|---|---|---|---|---|---|
| *Arc* | | | | | | | |
| | **Age x Sex** | 0.007 | 0.002 | 0.003 to 0.01 | 3.28 | 0.001 | * |
| | Age x Stress | 0.002 | 0.002 | -0.002 to 0.007 | 1.04 | 0.30 | ns |
| | **Sex x Stress** | -0.22 | 0.09 | -0.41 to -0.04 | 2.40 | 0.02 | * |
| *Egr1* | | | | | | | |
| | Age x Sex | -0.0001 | 0.004 | -0.008 to 0.007 | 0.027 | 0.98 | ns |
| | **Age x Stress** | -0.008 | 0.004 | -0.01 to -1.489e-005 | 1.99 | 0.0496 | * |
| | Sex x Stress | -0.05 | 0.16 | -0.36 to 0.26 | 0.32 | 0.75 | ns |
| *Fos* | | | | | | | |
| | Age x Sex | -0.005 | 0.004 | -0.01 to 0.002 | 1.36 | 0.18 | ns |
| | **Age x Stress** | -0.02 | 0.004 | -0.03 to -0.01 | 4.88 | <0.0001 | * |
| | Sex x Stress | 0.21 | 0.16 | -0.10 to 0.52 | 1.33 | 0.19 | ns |
| *Fosb* | | | | | | | |
| | **Age x Sex** | -0.01 | 0.005 | -0.02 to -0.001 | 2.18 | 0.03 | * |
| | **Age x Stress** | -0.03 | 0.005 | -0.04 to -0.01 | 4.82 | <0.0001 | * |
| | Sex x Stress | 0.33 | 0.21 | -0.09 to 0.76 | 1.55 | 0.12 | ns |
| *Junb* | | | | | | | |
| | Age x Sex | 0.007 | 0.004 | -0.002 to 0.01 | 1.49 | 0.14 | ns |
| | Age x Stress | -0.007 | 0.005 | -0.02 to 0.002 | 1.49 | 0.14 | ns |
| | Sex x Stress | -0.14 | 0.19 | -0.51 to 0.23 | 0.76 | 0.45 | ns |
| *Jund* | | | | | | | |
| | **Age** | -0.03 | 0.007 | -0.04 to -0.01 | 3.79 | 0.0003 | * |
| | Sex | 0.15 | 0.08 | -0.007 to 0.30 | 1.89 | 0.06 | ns |
| | Stress | -0.03 | 0.08 | -0.18 to 0.13 | 0.34 | 0.74 | ns |

*Outcome: gene expression relative to Control PN21Female; ns = not significant; *p < 0.05; Sex (Female, Male); Stress (Control, CVS)*

Table S11. Full statistical reporting from 3-way ANOVA analysis in Experiment 2

| Gene | Factor | F value | P value | Significance |
|---|---|---|---|---|
| *Arc* | | | | |
| | Age | F (1, 65) = 1.11 | P=0.30 | ns |
| | Sex | F (1, 65) = 2.33 | P=0.13 | ns |
| | Stress | F (1, 65) = 0.92 | P=0.34 | ns |
| | Age x Sex | F (1, 65) = 3.55 | P=0.06 | ns |
| | Age x Stress | F (1, 65) = 0.07 | P=0.79 | ns |
| | Sex x Stress | F (1, 65) = 2.16 | P=0.15 | ns |
| | Age x Sex x Stress | F (1, 65) = 3.38 | P=0.07 | ns |
| *Egr1* | | | | |
| | **Age** | F (1, 66) = 4.71 | P=0.03 | * |
| | Sex | F (1, 66) = 0.03 | P=0.87 | ns |
| | **Stress** | F (1, 66) = 7.12 | P=0.009 | * |
| | Age x Sex | F (1, 66) = 0.02 | P=0.89 | ns |
| | Age x Stress | F (1, 66) = 0.12 | P=0.72 | ns |
| | Sex x Stress | F (1, 66) = 1.14 | P=0.29 | ns |
| | **Age x Sex x Stress** | F (1, 66) = 4.27 | P=0.04 | * |
| *Fos* | | | | |
| | Age | F (1, 61) = 1.25 | P=0.27 | ns |
| | Sex | F (1, 61) = 0.01 | P=0.90 | ns |
| | **Stress** | F (1, 61) = 20.71 | P<0.0001 | * |
| | Age x Sex | F (1, 61) = 0.003 | P=0.95 | ns |
| | Age x Stress | F (1, 61) = 0.008 | P=0.93 | ns |
| | Sex x Stress | F (1, 61) = 0.08 | P=0.78 | ns |
| | Age x Sex x Stress | F (1, 61) = 2.46 | P=0.12 | ns |
| *Fosb* | | | | |
| | Age | F (1, 61) = 0.04 | P=0.83 | ns |
| | **Sex** | F (1, 61) = 4.10 | P=0.047 | * |
| | **Stress** | F (1, 61) = 31.37 | P<0.0001 | * |
| | **Age x Sex** | F (1, 61) = 4.97 | P=0.03 | * |
| | Age x Stress | F (1, 61) = 0.04 | P=0.85 | ns |
| | Sex x Stress | F (1, 61) = 1.27 | P=0.26 | ns |
| | Age x Sex x Stress | F (1, 61) = 0.04 | P=0.84 | ns |
| *Junb* | | | | |
| | **Age** | F (1, 64) = 19.23 | P<0.0001 | * |
| | Sex | F (1, 64) = 2.82 | P=0.10 | ns |
| | **Stress** | F (1, 64) = 7.15 | P=0.009 | * |
| | **Age x Sex** | F (1, 64) = 5.58 | P=0.02 | * |
| | Age x Stress | F (1, 64) = 2.37 | P=0.13 | ns |
| | Sex x Stress | F (1, 64) = 0.46 | P=0.50 | ns |
| | Age x Sex x Stress | F (1, 64) = 1.52 | P=0.22 | ns |
| *Jund* | | | | |
| | **Age** | F (1, 66) = 32.96 | P<0.0001 | * |
| | **Sex** | F (1, 66) = 5.52 | P=0.02 | * |
| | Stress | F (1, 66) = 0.03 | P=0.86 | ns |
| | **Age x Sex** | F (1, 66) = 9.63 | P=0.003 | * |
| | Age x Stress | F (1, 66) = 1.29 | P=0.26 | ns |
| | Sex x Stress | F (1, 66) = 1.22 | P=0.27 | ns |
| | Age x Sex x Stress | F (1, 66) = 0.008 | P=0.93 | ns |

*Outcome: gene expression (relative to PN21 Females);*
*ns = not significant; *p < 0.05; Age (PN28, PN35);*
*Sex (Female, Male); Stress (Control, CVS)*

#### Supplemental Figures

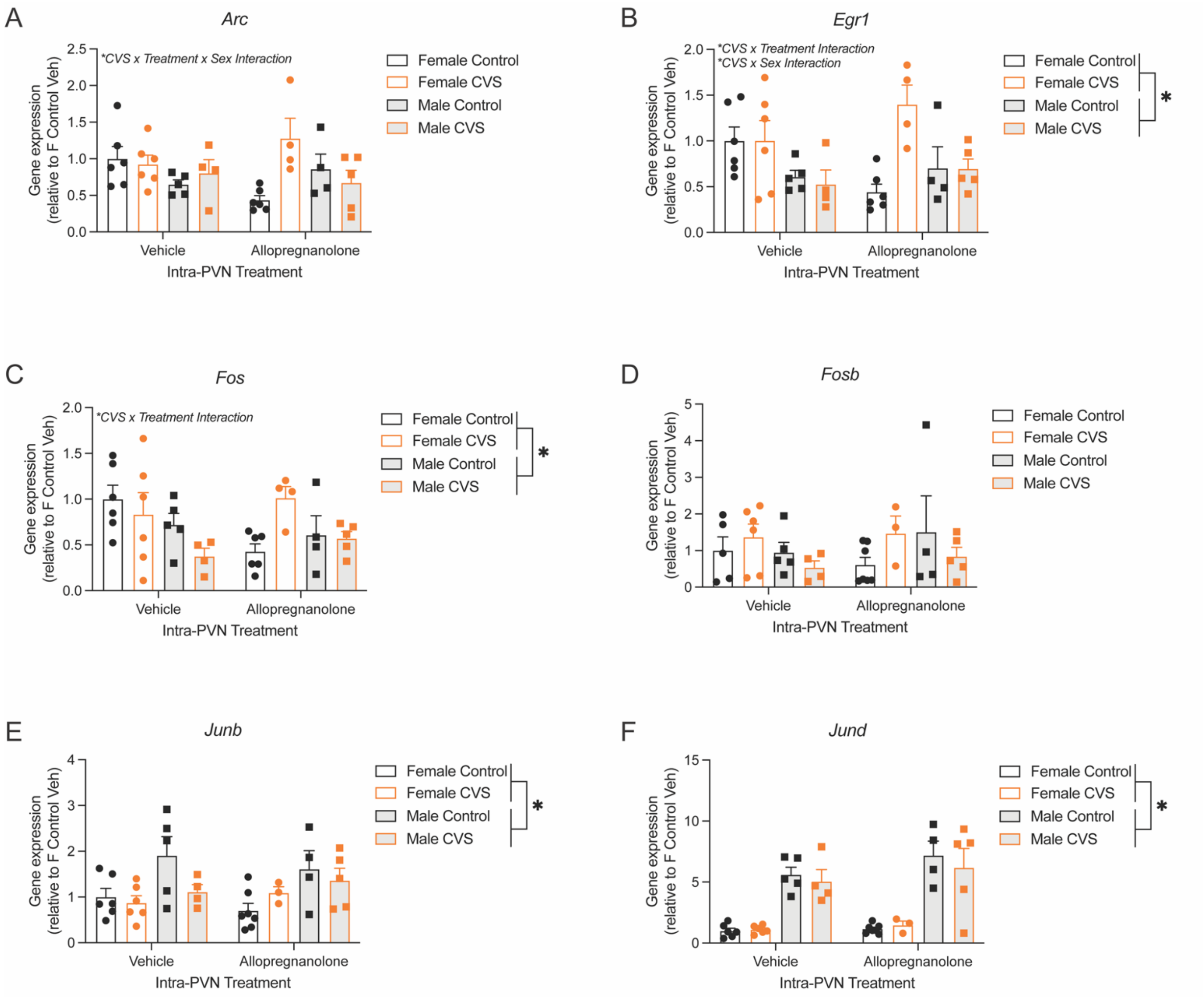


**Figure S1. Pubertal stress and intra-PVN allopregnanolone interacted to influence baseline gene expression.** Baseline gene expression in the PVN 2 hours after intra-PVN injection of vehicle or allopregnanolone for the 'puberty stress sensitive' immediate early genes (A) *Arc*, (B) *Egr1*, (C) *Fos*, (D) *Fosb*, (E) *Junb*, and (F) *Jund*. CVS = chronic variable stress; PVN = paraventricular nucleus. *$p < 0.05$ for main effect of sex or for specified interaction terms following 3-way ANOVA analysis. Data are mean ± SEM.

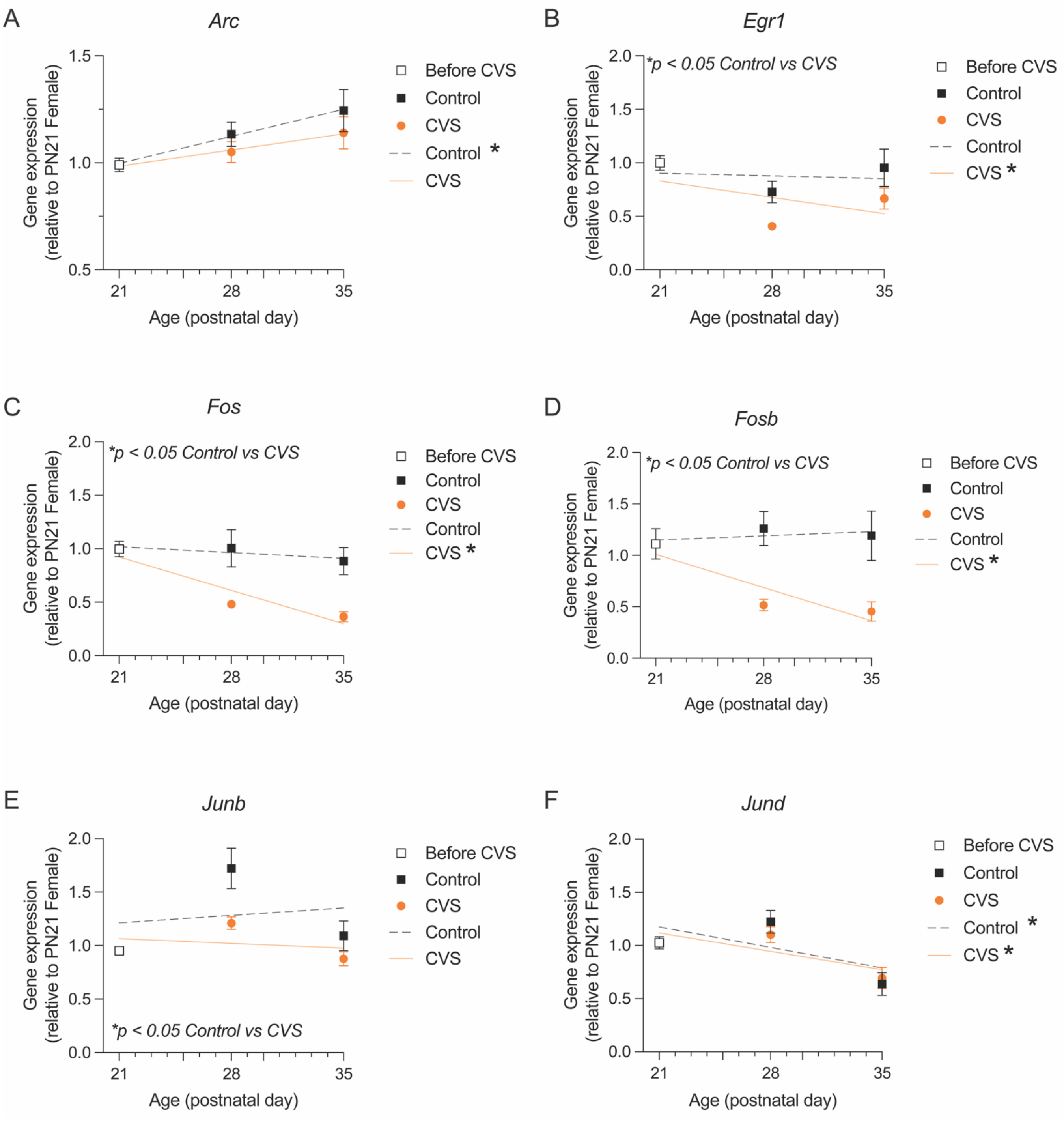


**Figure S2. Pubertal stress altered the trajectory of baseline IEG expression during adolescence.** Baseline gene expression in the PVN at PN21, PN28, and PN35 for the immediate early genes **(A)** *Arc*, **(B)** *Egr1*, **(C)** *Fos*, **(D)** *Fosb*, **(E)** *Junb*, and **(F)** *Jund*. At PN21, mice have not yet undergone pubertal CVS, so all PN28 and PN35 expression values were compared to PN21. For all genes, CVS exposure altered the trajectory of expression. For some genes (*Arc, Jund*), there was a non-zero trajectory of a change in gene expression during the early adolescent window in Control animals that was disrupted in animals exposed to pubertal CVS. For other genes (*Egr1, Fos, Fosb*), there was no significant difference in the gene expression in the PVN of Control animals from PN21-35. However, pubertal stress resulted in an alteration of this trajectory, decreasing gene expression during this time period. We further tested if the Control and CVS lines were different for each gene. For *Fos* and *Fosb*, the overall slopes of the trajectory in Control and CVS mice were significantly different. For *Egr1, Junb*, and *Jund*, the overall elevations, or intercepts, were significantly different between Control and CVS mice. CVS = chronic variable stress; IEG = immediate early gene. *$p < 0.05$ of linear regression, indicating slope is significantly non-zero, or for indicated differences between the Control and CVS linear regression lines. Data are mean ± SEM.

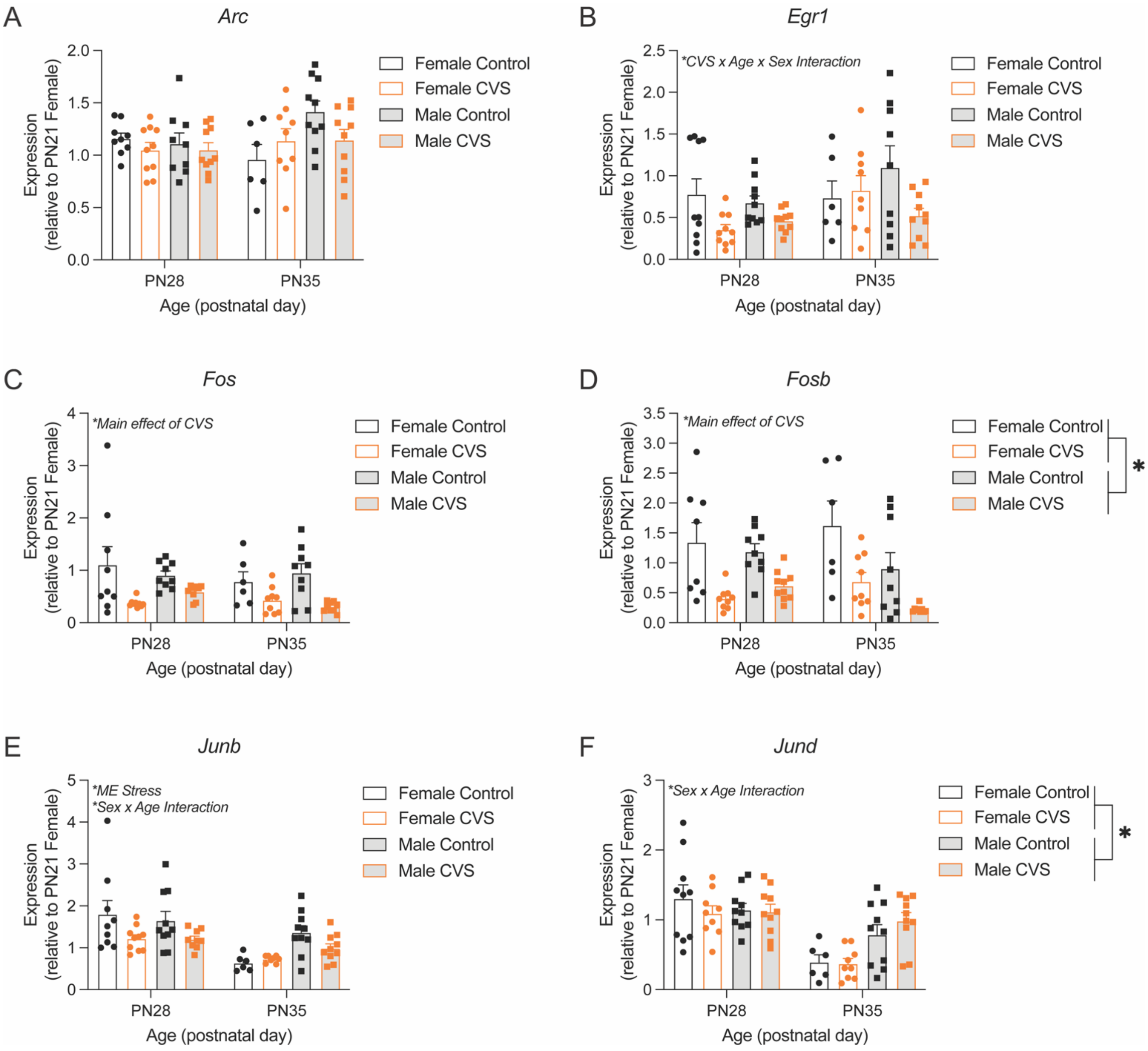


**Figure S3. Pubertal stress, sex, and age interacted to influence baseline IEG expression during adolescence.** Baseline gene expression in the PVN at PN28 and PN35 for the 'puberty stress sensitive' immediate early genes (A) *Arc*, (B) *Egr1*, (C) *Fos*, (D) *Fosb*, (E) *Junb*, and (F) *Jund*. (A) There was no effect of any factor on expression of *Arc* in this analysis. (B) For *Egr1*, there was a 3-way interaction between pubertal CVS, age of expression measurement, and sex of the mouse. While there was an overall increase in *Egr1* expression at PN35, which was driven by the finding that *Egr1* expression is blunted in male PN35 CVS relative to male PN35 Control mice. (C) *Fos* expression was uniformly influenced by CVS in females and males at both PN28 and PN35, such that there was decreased expression in CVS mice compared to Control mice. (D) There was a similar main effect of CVS on *Fosb* expression, such that there was decreased expression in pubertally stressed mice. Additionally, there was a main effect of sex, such that females had higher relative expression than males. (E) The expression of *Junb* was also significantly blunted in mice that had been exposed to pubertal CVS. *Junb* expression was influence by a sex by age interaction, where females had decreased expression at PN35 regardless of CVS exposure. (F) There was no influence of pubertal stress on *Jund* in this analysis. There was a sex by age interaction like *Junb*, such that females had decreased expression at PN35 regardless of CVS exposure. CVS = chronic variable stress; IEG = immediate early gene; PVN = paraventricular nucleus. *$p < 0.05$ for main effect of sex or for specified interaction term following 3-way ANOVA analysis. Data are mean ± SEM.
